# The unique lipopolysaccharide composition of *Salmonella* Typhimurium ST313 dampens pyroptosis and inflammasome activation and suppresses host cell death

**DOI:** 10.1101/2025.09.29.679141

**Authors:** Gabrielle Lê-Bury, Chantal Deschamps, Floriane Herit, Alexander V. Predeus, Audrey Dumas, Melita A. Gordon, Jay C. Hinton, Petr Broz, Florence Niedergang

## Abstract

In humans, non-typhoidal *Salmonella* (NTS) serovars, such as *Salmonella enterica* Typhimurium cause self-limiting gastroenteritis. However, invasive non-typhoidal *Salmonella* (iNTS) isolates ST313 have emerged in sub-Saharan Africa that cause a severe bacteremia in immunocompromised individuals including HIV-positive adults. Despite a decade of studies comparing ST313 to the classical diarrheal *S.* Typhimurium ST19 pathovariant, the molecular basis of the differences of pathogenesis between these bacteria remains unclear. Here we used primary human macrophages infected with the invasive *S.* Typhimurium ST313 D23580 isolate and the reference ST19 4/74 isolate as an *in vitro* model of infection. Transcriptomic profiling revealed that primary human macrophages infected with ST313 bacteria had distinct patterns of host cell death and inflammation responses compared to ST19-infected cells. Specifically, infection with ST313 isolate induced significantly less pyroptotic cell death which was associated with minimal processing of Gasdermin-D. We discovered that *S.* Typhimurium ST313 stimulated reduced IL-1β release and a low-level activation of the upstream inflammasome signaling cascade with reduced Caspase-4 and Caspase-1 cleavage. To investigate the molecular basis of this phenotype we purified lipopolysaccharide (LPS) species from ST313 to identify physical properties and structures that were distinct from ST19 LPS. The ST313-derived LPS was less pro-inflammatory than LPS from ST19, mirroring the low inflammatory phenotype of the live bacteria. Together, our data reveal that the unique LPS of ST313 is responsible for the reduced level of pyroptosis in human macrophages. This phenotype is consistent with the intracellular persistence and dissemination of these pathogenic bacteria.

**AUTHOR SUMMARY:** Non-typhoidal *Salmonella* (NTS) is a foodborne microbial pathogen that causes a self-limiting gastroenteritis in humans. However, in recent decades invasive non-typhoidal *Salmonella* (iNTS) isolates have emerged in sub-Saharan Africa, and are currently causing severe bloodstream infections responsible for about 66,000 deaths each year, largely in immunocompromised individuals. To investigate differences in pathogenicity between iNTS and NTS strains, we studied the infection of primary human macrophage with an iNTS ST313 isolate from Malawi in comparison to a gastroenteritis-associated ST19 reference strain. Primary macrophages play a key role in clearing *Salmonella* infections by recognizing bacterial components such as lipopolysaccharide (LPS). LPS induces immune responses including activation of cytokine release, inflammation and host cell death for effective bacterial elimination. Our study revealed that the iNTS isolate survived better in human primary macrophages than the NTS strain due to reduced inflammatory cell death, weakened inflammasome activation and lower cytokine release. We suggest that the distinct LPS structure is responsible for these dampened immune responses which aid the spread and persistence of *S.* Typhimurium ST313 in the human host.

## INTRODUCTION

*Salmonella enterica* (*S. enterica*) is a foodborne pathogen responsible for widespread disease in humans and other animals. Based on clinical manifestations in humans, bacterial serovars are classified into two groups: typhoidal and nontyphoidal (NTS). The typhoidal serovars *S.* Typhi and *S.* Paratyphi are restricted to humans and cause typhoid fever [1, 2]. In contrast, NTS serovars such as *S.* Enteritidis and *S.* Typhimurium typically cause self-limiting gastroenteritis in humans [3]. However, in recent decades invasive non-typhoidal *Salmonella* (iNTS) serovars have emerged that are causing an invasive disease and bloodstream infections resulting in the deaths of about 66,000 people each year in sub-Saharan Africa [4, 5]. Most cases are associated with malaria, malnutrition or an immunocompromised status [6–8]. Phylogenetic analysis of *S.* Typhimurium isolates that cause systemic disease in sub-Saharan Africa have identified a predominant multidrug-resistant multi-locus sequence type, ST313 [9, 10]. The emergence of the iNTS lineages was associated with a distinct genetic signature of ST313 and associated with altered pathogenicity, fitness and host adaptation [11–16]. The iNTS pathovariants are capable of hyper-dissemination and bacterial colonization of systemic sites in different animal models [17–20]. Several studies show that ST313 survive better than gastroenteritis-associated isolate ST19 in murine and human macrophages [19–23]. In human cell lines and primary mouse macrophages, the enhanced intracellular survival by ST313 is explained by a reduced induction of cytotoxicity compared with ST19 [20, 21].

*Salmonella* can trigger different types of cell death including a pro-inflammatory cell death called pyroptosis, a pro-inflammatory pathway originally defined as a Caspase-1- dependent process linked to the release of IL-1β and IL-18 [24–27]. Cytokine release and cell swelling are believed to occur upon the formation of pores generated by the assembly of cleaved Gasdermin-D (GSDMD), with GSDMD and IL-1β being cleaved by activated Caspase-1 [28–31]. Pyroptosis, which is necessary for the elimination of microbes, is typically initiated by inflammasome activation after pathogen recognition by the cells [26, 27, 31].

The current study builds on previous research that identified reduced inflammasome signaling and lower levels of cell death induction after ST313 infection compared to ST19 infection [20, 21]. Using human primary macrophages, we demonstrate that the difference of bacterial survival between ST313 D23580 and ST19 4/74 is likely driven by limited activation of the non-canonical inflammasome *via* Caspase-4, leading to decreased pyroptosis following ST313 infection. Our findings suggest that the enhanced capacity of ST313 to evade macrophage detection and stimulate low levels of inflammatory signaling is associated with previously uncharacterized differences in the structure and properties of LPS of this iNTS pathovariant.

## RESULTS

### ST313 bacteria trigger regulation of apoptotic process and inflammatory pathways in primary human macrophages

We have previously shown, with a gentamicin protection assay, that iNTS ST313 D23580 survives better within human monocyte-derived macrophages (hMDMs) than NTS ST19 4/74, without a corresponding difference in bacterial uptake [22]. This phenotype could be supported by a more favorable bacterial replication within the cells. Therefore, we analyzed the number of bacteria per cell using fluorescent microscopy (Fig 1A). We noticed that only 20% of ST313- infected macrophages had a high bacterial load (>10 per cell), while ST19-infected macrophages with a high burden represent 47.32% (Fig 1B). This indicates that ST313 do not replicate better than ST19 in primary human macrophages.

**Figure 1.**
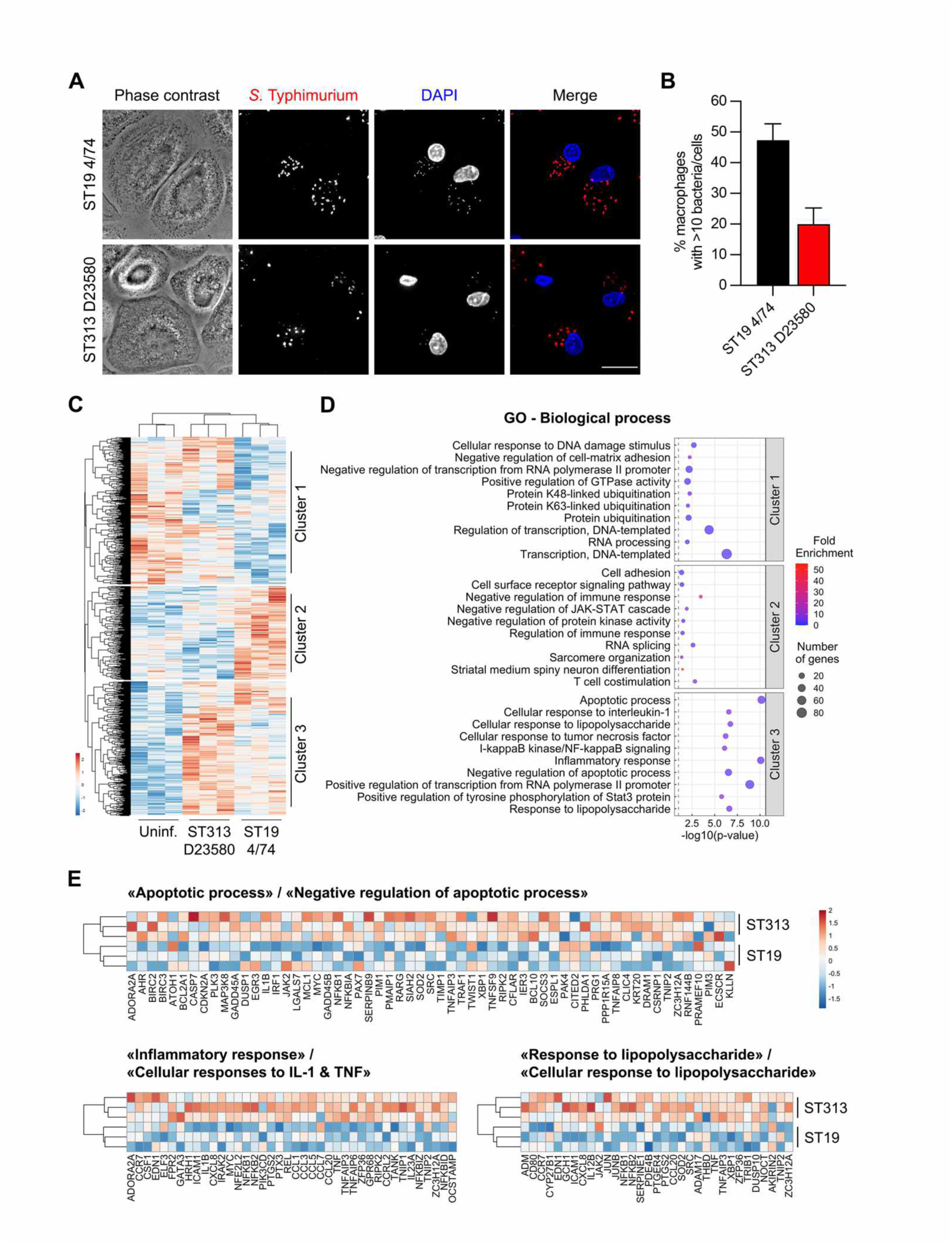
ST313 bacteria trigger regulation of apoptotic process and inflammatory pathways in primary human macrophages. (**A**) Human monocyte-derived macrophages (hMDMs) were infected with *Salmonella* Typhimurium ST19 4/74 (upper panel) or ST313 D23580 (lower panel) for 6h at 37°C. Then cells were fixed and labeled with anti-LPS antibody (second column in red) and DAPI (third column in blue). Scale bar, 20 μm. (**B**) Quantification of (A) representing the percentage of cells containing more than 10 bacteria for ST19 4/74 (black) and ST313 D23580 (red). Mean and SEM are represented for 3 different donors. (**C-E**) hMDMs were infected or not with ST19 4/74 or ST313 D23580 for 2h at 37°C. (**C**) Heatmap analysis of microarray data showing all genes differentially expressed in macrophages using hierarchical clustering. Red and blue colors indicate differentially up- and downregulated genes, respectively. *k*-means clustering on 3 donors. (**D**) Gene Ontology (GO) analysis of the top 10 of the most representative and significant Biological Processes (BP) from Cluster 1, Cluster 2 and Cluster 3 identified in (C). The x-axis represents -log10(*p*-value), dot size and dot color indicate the number of genes and the Fold Enrichment associated with the process, respectively. (**E**) Heatmap analysis of microarray data showing hierarchical clustering using *k*-means (on 3 donors) of genes involved in “Apoptotic process/Negative regulation of apoptotic process” (top heatmap), “Inflammatory response/Cellular responses to IL-1 & TNF” (bottom left heatmap) and “Response to lipopolysaccharide/Cellular response to lipopolysaccharide” (bottom right heatmap) from hMDMs infected with ST19 4/74 (bottom rows) or ST313 D23580 (top rows).

To understand why ST313 survive better in these cells [22], we analyzed global gene expression profiles of infected or uninfected hMDMs with each bacterial strain for 2h (Fig 1C-E). The ST19- and ST313-infected macrophages showed similar gene expression profiles that differed from the non-infected macrophages (Fig 1C). However, the gene expression profiles suggested that macrophages had responded differently to the two bacterial strains, revealing differential intracellular sensing by hMDMs (Fig 1C-D). Interestingly, Gene Ontology analysis of Cluster 3 showed an enrichment in pathways related to cell death, inflammation and response to lipopolysaccharide (LPS) (Fig 1D). The negative regulation of apoptotic processes and the inflammatory response signatures were up-regulated in macrophages infected with ST313 D23580 compared to those infected with ST19 4/74 (Fig 1E). Our results indicated that intrinsic properties of ST19 D23580 and ST313 4/74 are differentially-sensed by hMDMs, involving cell death, inflammatory and LPS responses. We suggest that the differences between the microscopy results (Fig 1A-B) and gentamicin protection assays [22] with hMDMs could be explained by a differential bacterial-induced cell death stimulated by ST313 or ST19 infection.

### ST313 D23580 induces less pyroptosis than ST19 4/74 in primary human macrophages

We next investigated the cytotoxic impact of *Salmonella* infection on hMDMs with the Lactate Dehydrogenase (LDH) assay, and observed less LDH release in the supernatant for ST313 D23580 compared to ST19 4/74 infection (Fig 2A). Our findings suggests that ST313 D23580 induces less cytotoxicity than ST19 4/74, consistent with the previous studies in other macrophage models [20, 21]. We then monitored the incorporation of propidium iodide (PI) in bacteria-infected hMDMs in real time (Fig 2B-C). PI incorporation was slower inside the cells infected with ST313 D23580 than with ST19 4/74, indicating a slower loss of membrane permeability in the ST313-infected macrophages. In addition, we observed cell swelling overtime after bacterial infection (Fig 2D), which is characteristic of pyroptosis, an inflammatory form of programmed cell death [32]. Pyroptosis is also defined by GSDMD pore formation at the plasma membrane and IL-1β release in the supernatant [28–31]. The pore formation is due to oligomerization of cleaved N-terminal domain of GSDMD after processing, which can be analyzed by Western blot. In the short timeframe after bacterial infection, we observed that GSDMD cleavage is less induced by ST313 D23580 than ST19 4/74 (Fig 2E). Furthermore, ST313 D23580 led to a reduced cleavage and release of IL-1β in the supernatant compared to ST19 4/74 (Fig 2F). Our findings showed that ST313 D23580 induced less pyroptosis than ST19 4/74 in primary human macrophages.

**Figure 2.**
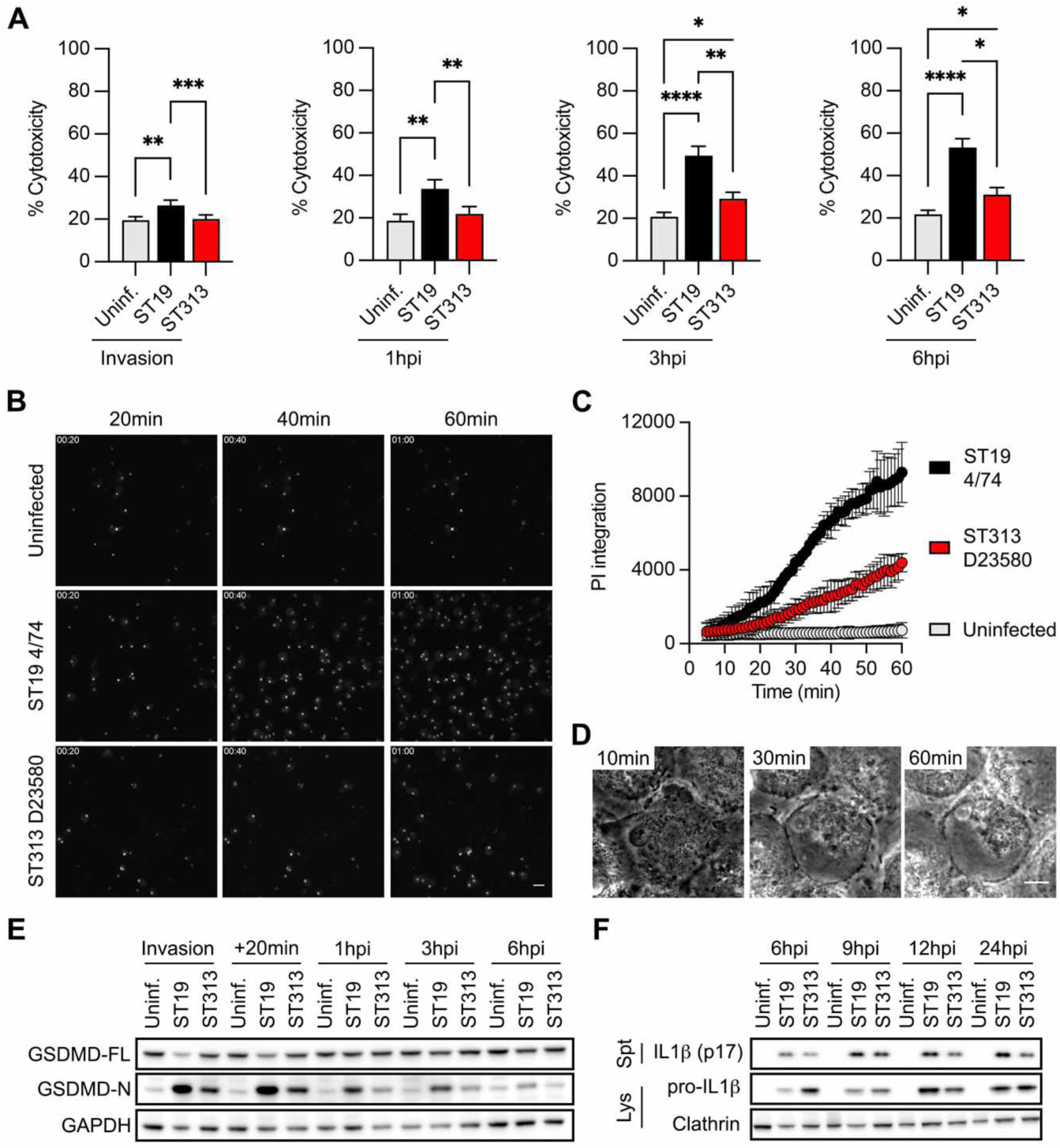
ST313 D23580 induces less pyroptosis than ST19 4/74 in primary human macrophages. Human monocyte-derived macrophages (hMDMs) were not infected (grey) or infected with *Salmonella* Typhimurium ST19 4/74 (black) or ST313 D23580 (red) during the indicated times. (**A**) Cytotoxicity was quantified using LDH assay at 20min after contact (“Invasion”, left); 1h; 3h and 6h post-invasion (hpi) and gentamicin treatment (middle and right graphs). The results are represented as a percentage of LDH release in the supernatants related to the total amount of LDH after total cell lysis in a control non-treated well. Friedman test was used (*, p-value<0.03 ; **, p-value<0.02 ; ***, p-value<0.0002 ; ****, p-value<0.0001) on 14 (“Invasion”) ; 7 (“1hpi”) ; 16 (“3hpi”) and 10 (“6hpi”) independent donors. (**B**) Cells were incubated with propidium iodide (PI) at 16µg/ml and infected or not (top panel) with *S.* Typhimurium ST19 4/74 (middle panel) or ST313 D23580 (bottom panel) for 1h. Images were acquired every minute with a spinning disk confocal microscope equipped with 5% CO2 and a heated chamber at 37°C. (**D**) Quantification from (C) are expressed as the total area of the PI fluorescent signal per field (in square pixels) overtime on 2 independent donors with biological duplicate. (**D**) Zoom of phase contrast in a movie showing a cell swelling upon infection with S. Typhimurium ST19 4/74. Scale bar, 10µm. (**E**) Western blot analysis of total lysates from hMDMs infected (or not) with ST19 4/74 or ST313 D23580 at different time points post-invasion (hpi), with anti-GSDMD antibody, and GAPDH as loading control. (**F**) Western blot analysis of total lysates and supernatants from hMDMs infected (or not) with ST19 4/74 or ST313 D23580 at different time points post-invasion (hpi), with anti-IL-1β antibody, and Clathrin as loading control.

### ST313 D23580 stimulates attenuated inflammasome activation compared to ST19 4/74 in primary human macrophages

GSDMD and IL-1β can be cleaved by Caspase-1, which itself is activated by proteolysis (Fig 3A). After 30 min of bacterial invasion, early Caspase-1 processing was observed. However, the extent of this cleavage was lower with ST313 D23580 than ST19 4/74 (Fig 3B). In addition to Caspase-1, IL-1β production was also regulated by Caspase-8 proteolysis and recruitment to the inflammasome along with Caspase-1, NLRP3 and NLRC4 (Fig 3A). Similar to the Caspase-1 and GSDMD findings, the Caspase-8 processing was strongly attenuated after ST313 infection compared to ST19 infection (Fig 3C).

**Figure 3.**
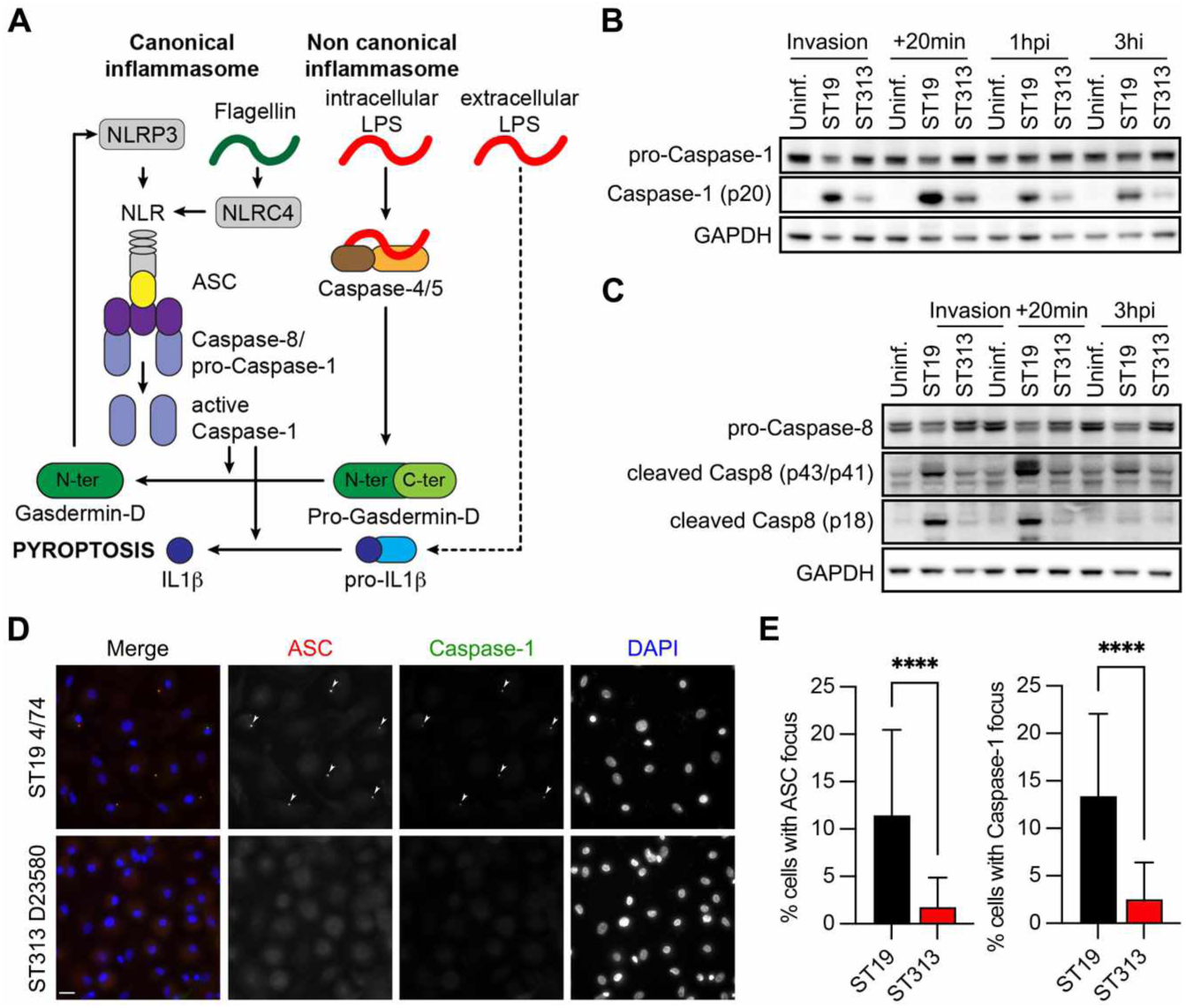
ST313 D23580 stimulates attenuated inflammasome activation than ST19 4/74 in primary human macrophages. (**A**) Schematic illustration of inflammasome activation in human macrophages upon exposure to LPS or flagellin of *Salmonella* Typhimurium (35, 36, 38). Flagellin activates the canonical inflammasome NLRC4 activation and oligomerization of NLRC4/NLRP3/ASC/pro-Caspase-8/pro-Caspase-1, leading to Caspase-1 processing, IL-1β and GSDMD cleavage, then pyroptosis. Intracellular LPS activated the non-canonical inflammasome leading to complex LPS/Caspase-4/5 formation that induces GSDMD cleavage and pyroptosis, including NLRP3 inflammasome activation as well. Extracellular LPS promotes transcription of pro-IL-1β. (**B-C**) Western blot analysis in total lysates of hMDMs infected (or not) with ST19 4/74 or ST313 D23580 at different time points post-invasion (hpi), with anti–Caspase-1 (**B**) or anti-Caspase-8 (**C**) antibodies, and GAPDH as loading control. (**D**) Representative images of hMDMs infected with ST19 4/74 (upper panel) or ST313 D23580 (lower panel) for 20min, stained for ASC (red, second column), Caspase-1 (green, third column), and DAPI for the nuclei (blue, last column). Arrowheads indicate ASC and Caspase-1 foci and Z-stacks images were acquired with Leica widefield microscope and maximum projection images are shown. Scale bar, 20µm. (**E**) The percentage of cells with ASC focus (left) or Caspase-1 focus (right) were calculated on three different donors with 25 images per donor and *t*-test analysis was used (****, p-value<0.0001).

Caspase-1 becomes activated through the canonical inflammasome pathway (Fig 3A), where pro-Caspase-1 is recruited to the inflammasome signaling platform *via* the ASC adaptor protein [33]. Activation of the inflammasome can be visualized through detection of Caspase-1 and ASC-speck formation by immunofluorescence (Fig 3D). For both proteins, fewer macrophages displayed fluorescent foci after ST313 infection compared to ST19 infection, indicating reduced canonical inflammasome activation by the iNTS strain (Fig 3D-E).

Overall, these results show that the invasive *S.* Typhimurium ST313 D23580 triggers less canonical inflammasome activation than the gastroenteritis-associated ST19 bacteria in primary human macrophages.

### ST313 D23580 LPS triggered weaker non-canonical inflammasome activation and pyroptosis than ST19 4/74 LPS in primary macrophages

*Salmonella* Typhimurium ST19 induces the formation of non-canonical inflammasomes in human macrophages via Caspase-4-mediated LPS detection, to activate GSDMD and induce pyroptosis (Fig 3A) [34–37]. In hMDMs, we noticed that Caspase-4 was less processed after ST313 infection than ST19 infection, suggesting a role for LPS in the differing level of pyroptosis induction between the two *Salmonella* strains (Fig 4A). Micoli *et al.* have shown that the O-polysaccharide chain differs between ST313 D23580 and other *S.* Typhimurium strains [38]. Furthermore, Pulford *et al.* discovered that the *lpxO* gene of *S.* Typhimurium ST313 is a pseudogene due to a premature stop codon (E198*), leading to a lack of this LPS hydroxylation in ST313 [16].

**Figure 4.**
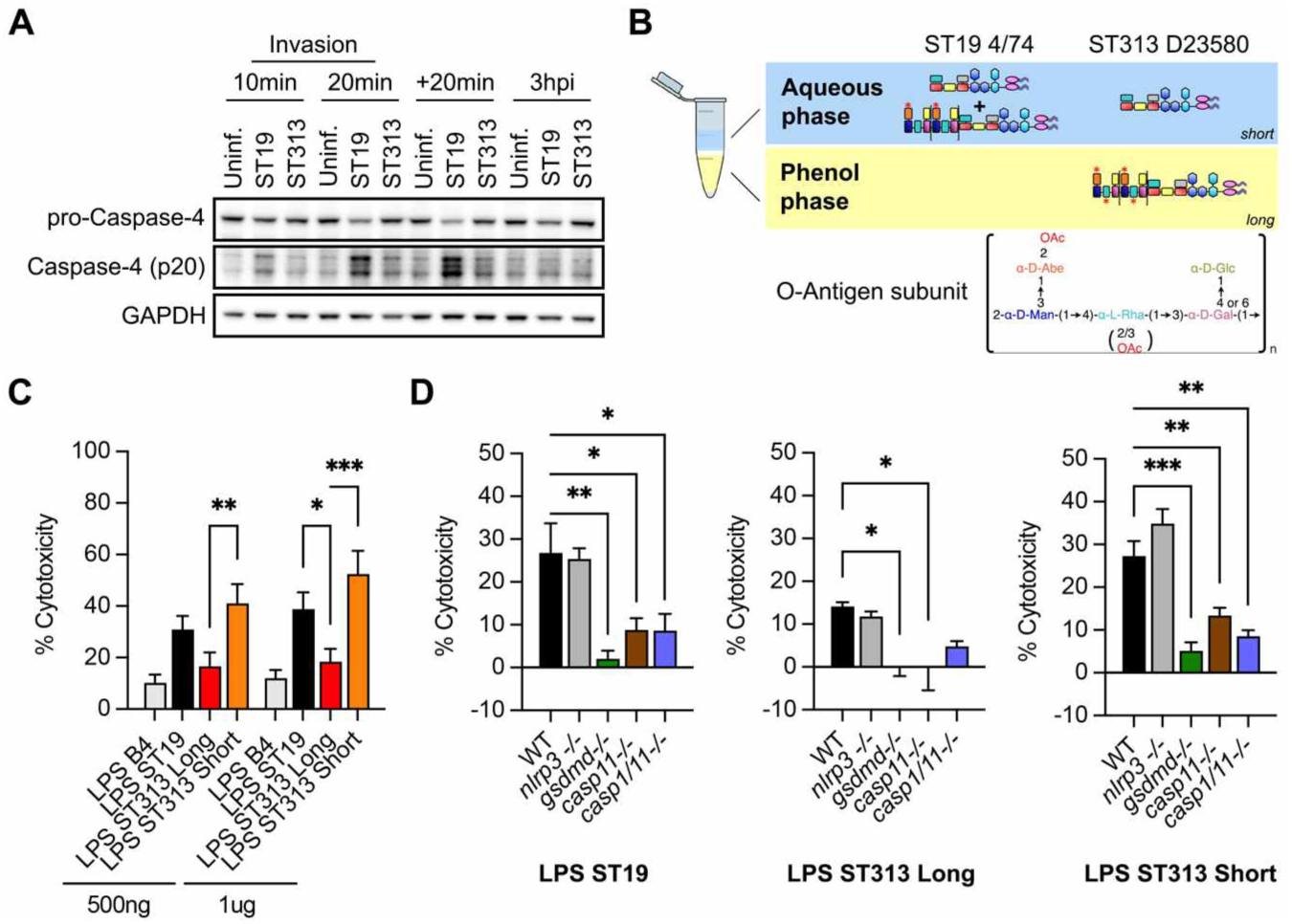
LPS from ST313 D23580 induce less pyroptosis through non-canonical inflammasome than LPS from ST19 4/74 in primary macrophages. (**A**) Western blot analysis of total lysates from hMDMs infected (or not) with ST19 4/74 or ST313 D23580 at different time points post-invasion (hpi), with anti-Caspase-4 antibody and GAPDH as loading control. (**B**) Schematic illustration of LPS purified from ST19 4/74 and ST313 D23580 (upper panel). Saccharides are pink ovals for the lipidA, Inner Core, Outer Core and O-antigen subunit are formed of blue hexagons, colorful horizontal rectangles, and colorful vertical rectangles, respectively. Red stars represent O-acetylation. Lower panel is the structure of repeating subunit of O-polysaccharide. O-acetylation in curved brackets varies between ST19 4/74 and ST313 D23580. (**C**) Primary bone marrow-derived macrophages (BMDMs) from WT mice were primed with LPS, then transfected with 500ng or 1μg of LPS B4, LPS ST19 4/74, LPS ST313 D23580 long or short. Cytotoxicity was measured with LDH assay in triplicate 2h post transfection. The mean of three independent experiments is shown and Two-way ANOVA was used (*, p-value<0.03 ; **, p-value<0.02 ; ***, p-value<0.0002). (**D**) BMDMs from WT, *nlrp3^-/^*^-^, *gsdmd^-/-^*, *casp11^-/-^* and *casp1/11^-/-^* mice were primed with LPS, then transfected with 1μg of LPS 4/74 ST19 (left), LPS ST313 D23580 long (middle) or short (right). Cytotoxicity was measured with LDH assay in technical triplicate of one experiement, the mean was plotted and one-way ANOVA was used (*, p-value<0.03 ; **, p-value<0.02 ; ***, p-value<0.0002).

We next purified LPS from ST313 D23580 and ST19 4/74. All the LPS from ST19 4/74 was purified as expected in the water phase by the phenol/water method [39]. In contrast, the rough LPS (short) and smooth LPS (long) from ST313 D23580 were found in both the water phase and phenolic phase, respectively (Fig 4B). This finding clearly indicates difference in LPS structure and solubility between the LPS which are mediated by the O-acetylation (OAc) of Rhamnose a (Rha) residue present in the O-chain polysaccharides of the ST313 strain. The O-acetylation increases hydrophobicity due to the presence of an already O-acetylated Abequose residue. LPS from ST313 D23580 (Fig 4B), containing the O-acetylated Rhamnose became soluble in the Phenol phase of the LPS extraction, as described [38]. An alternative purification method was then used to purify all LPS species from ST313 (D23580) [40].

To assess the impact of LPS on pyroptosis, we analyzed the cytotoxicity of the various fractions named LPS ST19, LPS ST313-long and LPS ST313-short, compared with the commercial *E. coli*-derived LPS BP4, in primary bone marrow-derived macrophages (BMDMs). We first induced pyroptosis by transfecting different amounts of the purified LPS variants into primed BMDMs from wild type (WT) mice. The results showed that transfection with LPS ST313-long induced less cytotoxicity than LPS ST19 or LPS ST313-short (Fig 4C).

To identify the pathways involved, cytotoxicity was compared between LPS-transfected BMDMs from WT mice or mice deficient for NLRP3 (*nlrp3-/-*), GSDMD (*gsdmd-/-*), Caspase-1 and -11 (*casp1/11-/-*) or Caspase-11 (*casp11-/-*). Caspase-11 is the murine homolog of human Caspase-4 (Fig 4D). These findings confirmed that pyroptosis triggered by the various LPS species proceeds to GSDMD activation *via* the non-canonical (Caspase-11) pathway, and was not dependent on the canonical NLRP3 pathway. The presence of the unique long LPS in ST313 apparently reduces the ability of ST313 bacteria to activate the non-canonical inflammasome pathway required for pyroptosis.

## DISCUSSION

In this study, we reveal that the unique LPS structure of ST313 prevents non-canonical inflammasome signaling and inhibits pyroptotic cell death in primary human macrophages. Our findings highlight the specific response of macrophages to these invasive bacteria and could explain dissemination of the pathogen *via* a “trojan horse” mechanism in host cells.

Under our experimental conditions, ST313 bacteria did not exhibit increased cellular invasiveness in gentamicin protection assays but showed a higher capacity for intracellular survival within human macrophages, compared with their ST19 counterparts [22]. These results align with and may explain the higher systemic colonization observed in animal models for ST313 relative to ST19 [17–21, 41–43]. Carden *et al.* demonstrated that ST313 hyper-disseminates from the gut to systemic sites in mice through CD11b+ dendritic cells, a process dependent on the loss of the SseI effector [42]. However, no differences were reported in survival between ST313 *versus* ST19 within murine dendritic cells *in vitro*. Similarly, our experiments revealed no differences in bacterial survival within CaCo2 epithelial cells (data not shown).

In contrast, human macrophages exhibited a remarkable capacity to distinguish between these ST313 and ST19 bacterial strains *in vitro*, shown by the distinct gene expression profiles we observed in infected primary human macrophages. Notably, the most striking differential cellular responses between the two bacterial strains were pathways associated with “inflammation” and “cell death”. Previous studies showed that ST313 bacteria were less cytotoxic than ST19 bacteria in a human cell line and in primary mouse macrophages [20, 21].

We used a combination of live-cell imaging and biochemical approaches to discover that this difference of cytotoxicity involves the pyroptotic inflammatory cell death which is characterized by Caspase-1 and GSDMD cleavage. This difference in cellular sensing could involve both the canonical inflammasome activated by flagellin, and the non-canonical inflammasome, triggered by cytosolic LPS [44, 45]. While flagellin might be involved [20], we demonstrate that this divergence may also reflect differences in the LPS structures of the two strains. We reveal that Caspase-4, the human homolog of murine Caspase-11 involved in the non-canonical inflammasome pathway [36, 46], was cleaved less in ST313-infected macrophages than ST19- infected macrophages. We conclude that the difference of pyroptosis induction between ST19 and ST313 in human macrophages depends on the differential sensing of bacterial components, including LPS.

In conclusion, human macrophages can distinguish between two closely-related bacterial pathovariants by activating distinct sensing mechanisms and gene expression patterns, reflecting differences in virulence. Our findings could pave the way for new strategies to better understand ST313, a pathogen characterized by human-associated niches, reservoirs or potential human to human transmission. Such insights could support better control strategies and help to reduce mortality in high-risk populations.

## MATERIALS AND METHODS

### Experimental model and study participant details

#### Primary human macrophages

Human monocytes-derived macrophages (hMDMs) are differentiated from peripheral blood mononuclear cells (PBMCs) from whole blood of healthy donors from Etablissement Français du Sang Ile-de-France, Site Trinité (Inserm agreement #15/EFS/012 and #18/EFS/030 ensuring that all donors gave a written informed consent and providing anonymized samples). The procedures for sample collection and conservation were declared and approved through CODECOH (COnservation d’Eléments du COrps Humain, No. DC-2021-4166) by the French Ministry of Higher Education, Research and Innovation, in accordance with French regulations on human biological elements. PBMCs are isolated by density gradient sedimentation using Ficoll-Plaque (GE Healthcare). This was followed by adhesion on plastic at 37°C for 2h in adhesion medium (RPMI 1640 (Life Technologies) supplemented with 100 μg/ml streptomycin/penicillin and 2mM L-glutamine (Invitrogen/Gibco)). Then, the adhered cells were washed once with warm adhesion medium and differentiated in complete macrophage medium (RPMI 1640 supplemented with 10% FCS (Eurobio), 100 μg/ml streptomycin/penicillin, 2 mM L-glutamine) and 10 ng/mL recombinant human macrophage colony-stimulating factor (rhM-CSF; R&D systems) [47, 48].

### Primary mouse macrophages

Bone marrow-derived macrophages were differentiated in DMEM (Gibco) supplemented with 20% MCSF (3T3 supernatant), 10% heat-inactivated FCS (Bioconcept), 10 mM HEPES (Bioconcept), 100 μg/ml penicillin/streptomycin (Bioconcept, 4-01H00-H) and non-essential amino acids (Gibco, 11140-035 at 1/100), and stimulated on day 7**–**9 of differentiation as in [49].

### Bacterial strains and growth conditions

*Salmonella enterica* Typhimurium ST313 D23580 and ST19 4/74 isolates were described [9, 50, 51]. *Salmonella enterica* Typhimurium SSS18, a version of *Salmonella enterica* Typhimurium ST313 D23580 that is Chloramphenicol and Ampicillin sensitive (kind gift from Robert Kingsley, Sanger Institute), was used to introduce the pMR98-DsRed replicative plasmid expressing DsRed protein [52]. Bacteria were grown with shaking in Lysogeny Broth (LB) at 37°C. Ampicillin was added at 50 μg/mL to grow recombinant bacteria. The absorbance at 600 nm of the bacterial suspensions was used to determine the multiplicity of infection (MOI), by estimating that 10^9^ bacteria per mL give an A600 of 1. The inoculum dose was then calculated by plating serial dilutions onto LB agar plates.

### Bacterial infection

*S.* Typhimurium were grown overnight at 37°C with shaking in Luria-Bertani broth, then subcultured for 2-3h with shaking in LB until late logarithmic phase was reached. The absorbance at 600 nm of the bacterial suspensions was used to determine the multiplicity of infection, by estimating that 10^9^ bacteria per mL give an A600 of 1. The bacterial suspension was diluted in RPMI without serum to have a calculated MOI of 30. The inoculum dose was calculated by plating serial dilutions onto LB agar plates. 1 mL of bacterial suspension inoculum was added to hMDMs in 6 well plates. Then, the 6-well plates were centrifuged for 2 minutes at 400 g to synchronize the infection and the kinetics was started by incubating the plates at 37°C, 5% CO2.

After a 20 minutes incubation at 37°C that corresponds to the contact step, hMDMs were gently washed 2 times with sterile PBS, and then incubated in complete RPMI medium supplemented with 100 µg/mL of gentamicin for the indicated times.

### Transcriptional profiling by Microarray

hMDMs of 3 different healthy donors, infected or not with HIV-1_ADA_WT, were co-infected with either *S*. Typhimurium D23580 or 4/74 for 2h. RNA was extracted using RNeasy® Plus mini kit (Qiagen) according to the manufacturer’s instructions. After validation of the RNA quality with Bioanalyzer 2100 (using Agilent RNA6000 nano chip kit), 50ng of total RNA were reverse transcribed following the Ovation PicoSL WTA System V2 (Nugen). Briefly, the resulting double strand cDNA was used for amplification based on SPIA technology. After purification according to Nugen protocol, 3.6ug of Sens Target DNA were fragmented and biotin labelled using Encore Biotin Module kit (Nugen). After control of fragmentation using Bioanalyzer 2100, cDNA was then hybridized to GeneChip® Human Gene 2.0 ST (Affymetrix) at 45°C for 17h. After overnight hybridization, chips were washed on the fluidic station FS450 following specific protocols (Affymetrix) and scanned using the GCS3000 7G. The scanned images were then analyzed with Expression Console software (Affymetrix) to obtain raw data (cel files) and metrics for Quality Controls. The observations of some of these metrics and the study of the distribution of raw data showed no outlier experiment. RMA normalization was performed using R with Version 17 of Entrezgene CDF brain array. Heatmaps are generated using R software with “pheatmap” function on normalized data, then clusters were extracted with “rect.hclust” function. Gene Ontology analysis (Biological Process) of extracted clusters were performed using DAVID Bioinformatics Resources [53, 54]. All data have been deposited in NCBI’s Gene Expression Omnibus and are accessible through GEO Series accession number GSE308645 (using the secure token ujufeiwwfpcjzyf).

### Cytotoxicity assay

Cytotoxicity induced by bacterial infection was measured using Lactate Dehydrogenase (LDH) release in the cell supernatant with the Cytotoxicity Assay Kit according to the manufacturer’s instructions (Pierce) for human macrophages and CytoTox 96 non-radioactive cytotoxicity assay (Promega) for mouse macrophages. Results are expressed as a percentage of total cellular LDH (100 % lysis).

### Live cell imaging of propidium iodide (PI) incorporation

hMDMs were detached using PBS/EDTA 2mM, and seeded on a μ-Slide 8 Well (Ibidi) at 1.5.10^5^ cells/well. Cells were infected at a MOI of 10 with 0.1 mL of bacterial suspension (*S*. Typhimurium ST313 D23580 or ST19 4/74). PI was added at the same time as bacteria at a final concentration of 160 μg/mL. Samples were observed with a spinning disk (CSU-X1M1; Yokogawa) confocal inverted microscope (DMI6000; Leica) equipped with a CoolSnap HQ^2^ camera (Photometrics) and a heated chamber at 37°C with 5% CO_2_ in a BSL3 laboratory. PI and PH images were acquired every 1 min for 1 h using a HCX PL FLUORTAR 10X/0.3 NA dry objective with the MetaMorph 7.5.5 software (Molecular Devices). Analyses were performed using custom-made ImageJ (National Institutes of Health) routines. To observe cell swelling, PH images were acquired every 30 s for 1h using a HCX PLAN APO 63X/1.4 NA oil PH objective.

### Immunoblot analysis

Whole cell lysates from hMDMs were obtained by addition of lysis buffer (Tris pH7.5 20 mM, NaCl 150 mM, NP40 0.5 %, NaF 50 mM, Orthovanadate 1 mM and Mini EDTA free Protease Inhibitor Mixture ; Roche). Cell lysates were separated by electrophoresis in denaturing 4-12 % gradient SDS-PAGE (Thermofisher), then transfer to PVDF membrane (Millipore). Membranes were incubated with diluted primary antibody in 5 % nonfat dry milk or 5 % BSA in 1X Tris Buffered Saline (TBS), 0.1 % Tween®20 at 4°C with gentle shaking, overnight: rabbit anti-IL1b (Abcam) in 5 % TBS with 0.1 % Tween®20 (TBST), rabbit anti-Caspase-1 (D7F10 ; Cell Signaling) in 5 % BSA TBST, rabbit anti-GSDMD (Atlas antibodies) in 5% nonfat dry milk TBST, rabbit anti-Caspase-4 (Cell Signaling), mouse anti-Caspase-8 (1C12 ; Cell Signaling) in 5% BSA TBST, rabbit anti-NLRC4 (D5Y8E ; Cell Signaling) in 5% BSA TBST, and as loading control, mouse anti-GAPDH (D4C6R ; Cell Signaling) 5% nonfat dry milk TBST, mouse anti-clathrin heavy chain (BD Transduction Laboratories) 5% nonfat dry milk TBST.

### LPS extraction, purification and characterization

LPS were extracted, purified and characterized at LPS-BioSciences (Paris-Saclay University, Orsay 91405, France, www.lpsbiosciences.com). Bacteria were first extracted by the classical phenol- water (PW) method [55] to produce ST19 and ST313 short, Rough-type LPS, isolated traditionally from the water-phase. LPS purification was performed as described previously by enzyme treatments and acidified solvent extraction for contaminants removal [56]. Alternatively, as the hydrophobic elements of the O-chain present in the ST313 LPS were soluble in the phenol-phase of the PW method, the ST313 complete LPS could be obtained by extraction with the ammonium hydroxide-isobutyric acid method [40], and compared to the short-chains ST313 LPS isolated from the water-phase of the PW method. Comparison of the two extraction methods showed the interest of the non-toxic, single phase ammonium hydroxide-isobutyric acid method that did not segregate LPS molecular species during extraction.

### Matrix Assisted Laser Desorption Mass spectrometry (MALDI-MS) analysis of lipid A samples

LPS samples were hydrolysed by the triethylamine-citric acid method and the lipid A extracts in chloroform-methanol-water as described [57]. MALDI-MS spectra were obtained as described [58].

### SDS-polyacrylamide gel electrophoresis of LPS

Gels were loaded with 0.2 to 0.5 g of R-type LPSs and with 1 to 2 g of Smooth-type LPSs. Both LPS-type preparations were electrophoresed as previously described [58] and then stained with silver nitrate.

The lipid A structure of both LPS were characterized and compared by MALDI-mass spectrometry. Both lipid A structures corresponded to a mixture of molecular species differing by the number of fatty acids. The major peaks corresponded in both cases to a hepta- and hexa-acylated molecular species. In both cases the Phosphate groups were decorated with small amounts of AraN, and the ST313 lipid A spectrum displayed in addition a small peak corresponding to an additional phosphate group (not shown).

### Inflammasome assay

Primary BMDMs were plated in 96-well plates at 5 x 10^4^ cells per well a day prior to stimulation. Cells were primed with ultrapure E. coli K12 LPS (InvivoGen) for 4 h in Opti-MEM and transfected using DOTAP (Sigma) with purified LPS at the concentrations indicated. Cytotoxicity was measured in the supernatant in triplicates 2h post transfection.

### Ethics statement

Inserm agreement #15/EFS/012 and #18/EFS/030 ensures that all blood donors gave a written informed consent and that EFS provides anonymized samples. The procedures for sample collection and conservation were declared and approved through CODECOH (COnservation d’Eléments du COrps Humain, No. DC-2021-4166) by the French Ministry of Higher Education, Research and Innovation, in accordance with French regulations on human biological elements.

### Quantification and statistical analysis

Student’s *t*-test , One- or Two-ANOVA using Prism software (GraphPad Software, San Diego, CA). A two-tailed *p-value* of <0.05 was taken to indicate statistical significance (* represents *p-value*<0.05, ** represents *p-value*<0.01, *** represents *p-value* < 0.001)

## ACKNOWLEDGMENTS

We thank EFS (Saint Vincent de Paul, Trinité and Saint-Antoine) for buffy coat supply, Sébastien Jacques and Florent Dumont (GENOM’IC facility of Institut Cochin) for initial transcriptomic analysis, the IMAG’IC facility of Institut Cochin that is part of the national France-BioImaging infrastructure supported by Agence Nationale de la Recherche (ANR-10-INBS-04), Simon Faillot for help on R software, and Florent Salvador for technical support. We thank Olivia Steele-Mortimer (NIH USA) and Serge Benichou (Institut Cochin) for initial discussions on the project. We thank also Thomas Henry (CIRI, Centre International de Recherche en Infectiologie) for discussions on inflammasomes and LPS Biosciences for their expert contribution on LPS purification.

## Funding

Work in the laboratory of FN is supported by CNRS, Inserm, Université Paris Cité ; Agence Nationale de Recherches sur le Sida et les Hépatites (ANRS AO2012-2 (FN, GLB, CD)) ; Fondation pour la Recherche Médicale (FRM DEQ20130326518 (FN, GLB)). This work was partly supported by the Wellcome Trust Investigator award 222528/Z/21/Z to J.C.D.H. For the purpose of open access, the authors have applied a CC BY public copyright licence to any Author Accepted Manuscript version arising from this submission.

## Author contributions

Conceptualization: F.N., C.D., G.L.-B.

Methodology: F.N., C.D., G.L.-B., J.C.H., M.A.G., P.B.

Formal Analysis: G.L.-B., C.D., A.V.P.

Investigation: G.L.-B., F.H., C.D., A.D., F.N.

Resources: F.N., J.C.H, M.A.G, P.B.

Writing – original draft: G.L.-B., F.N.

Writing – review & editing: G.L.-B., F.N, J.C.H.

Visualization: G.L.-B., F.N.

Project administration: F.N.

Supervision: F.N.

Funding acquisition: F.N.

## Declaration of interests

Authors declare that they have no competing interests.

## Notes

### Competing Interest Statement

The authors have declared no competing interest.

